# NanoMEA: a versatile platform for high-throughput analysis of structure-function relationships in human stem cell-derived excitable cells and tissues

**DOI:** 10.1101/453886

**Authors:** Alec S.T. Smith, Eunpyo Choi, Kevin Gray, Jesse Macadangdang, Eun Hyun Ahn, Elisa C. Clark, Phillip Tyler, Michael A. Laflamme, Leslie Tung, Joseph C. Wu, Charles E. Murry, Deok-Ho Kim

## Abstract

Somatic cells derived from human pluripotent stem cell (hPSC) sources hold significant potential as a means to improve current *in vitro* screening assays. However, their inconsistent ability to recapitulate the structural and functional characteristics of native cells has raised questions regarding their ability to accurately predict the functional behavior of human tissues when exposed to chemical or pathological insults. In addition, the lack of cytoskeletal organization within conventional culture platforms prevents analysis of how structural changes in human tissues affect functional performance. Using cation-permeable hydrogels, we describe the production of multiwell nanotopographically-patterned microelectrode arrays (nanoMEAs) for studying the effect of structural organization on hPSC-derived cardiomyocyte and neuronal function *in vitro*. We demonstrate that nanoscale topographic substrate cues promote the development of more ordered cardiac and neuronal monolayers while simultaneously enhancing cytoskeletal organization, protein expression patterns, and electrophysiological function in these cells. We then show that these phenotypic improvements act to alter the sensitivity of hPSC-derived cardiomyocytes to treatment with arrhythmogenic and conduction-blocking compounds that target structural features of the cardiomyocyte. Similarly, we demonstrate that neuron sensitivity to synaptic blockers is increased when cells are maintained on nanotopographically-patterned Nafion surfaces. The improved structural and functional capacity of hPSC-derived cardiomyocyte and neuronal populations maintained on nanoMEAs may have important implications for improving the predictive capabilities of cell-based electrophysiological assays used in preclinical screening applications.

---

The potential for human pluripotent stem cells (hPSCs) to revolutionize preclinical screening modalities has been widely discussed and, to some extent, already realized^1, 2, 3^. With regards to electrophysiological analysis, such cell sources enable assessment of patient-specific functional profiles to be developed at both the single cell and population level^4, 5^. The information gleaned from study of these cells has already improved our understanding of functional responses to pharmacological and pathological challenge in human tissues and will continue to do so as our ability to generate more specific cellular sub-types increases. However, hPSC-derived somatic cells are most commonly analyzed at fetal stages of development, which limits their ability to predict more mature human tissue behavior accurately^6^. Multiple methods for enhancing the maturation of hPSC-derived cells have been described previously^7, 8, 9, 10, 11, 12, 13, 14, 15^, and analysis of such engineered cells and tissues has demonstrated a capacity for matured human cells to exhibit responses to drug treatment or genetic perturbation that more closely recapitulate that of native adult human tissues^9, 10, 16^. Unfortunately, many of the maturation methods developed so far utilize unique platforms with configurations and/or analysis methods that can present difficulties when attempting to scale up to high-throughput formats. Methods to improve hPSC-derived somatic cell development that can be readily adapted to multiple cell lineages as well as existing high-throughput screening methodologies are therefore highly desirable.

To address this need, we present the development and validation of a nanotopographically-patterned, multiwell microelectrode array (MEA) as a means to enhance the structural organization and functional performance of hPSC-derived electrically-active cell types (cardiomyocytes and neurons) within an extant high-throughput electrophysiological assay. The device constitutes a simple and reproducible method utilizing bioinspired matrix nanotopographic cues to regulate structural development *in vitro* and thereby improve functional output. Data are presented that demonstrate the impact these surfaces have on hPSC-derived cardiomyocyte and neuron structural development as well as the effect these phenotypic changes have on both baseline and drug-induced electrophysiology in these cells. Specifically, we highlight the enhanced effect of compounds that target structurally-defined elements within the cardiac cell as well as drugs that inhibit synaptic connectivity in neuronal populations. Overall, the nanoMEA system represents the means to conduct in-depth analysis of how structural development and cytoskeletal organization within neuronal and cardiac monolayers affect functional output. In addition, this technology could enhance the capacity for preclinical MEA-based screens to predict drug efficacy and toxicity in humans.

## Results

### Integration of nanotopographically-patterned, ion-permeable thin films into microelectrode arrays

To facilitate structural organization of human cells within a hardware format well suited to high-throughput electrophysiological analysis, the following design criteria needed to be met: (1) a multiplexed system for sufficient throughput, (2) structural cues to promote cytoskeletal organization, (3) an integrated means of assessing electrophysiological activity in a non-invasive and longitudinal manner, and (4) optically transparent culture areas to facilitate assessment of cellular alignment prior to functional studies. To meet these criteria, we conceived of a method to integrate conductive, ion-permeable, nanotopographic patterns with 48-well MEA plates to promote hPSC-derived cardiomyocyte and neuron maturation while simultaneously enabling high-throughput assessment of population-level function (**Figure 1a**). Substrate topographic features in the high nm range have been shown to enhance the structural development of both hPSC-derived cardiomyocytes (hPSC-CMs) and neurons (hPSC-neurons)^17, 18, 19^. As such, 800 nm features (groove width_ridge width) were used throughout this study to create the nanoMEA devices. Pattern fidelity was assessed by scanning electron microscopy and the collected data validate the described methods in terms of their ability to imprint reproducible nanotopographies into the polymer substrate (**Figure 1b-d**).

**Figure 1:**
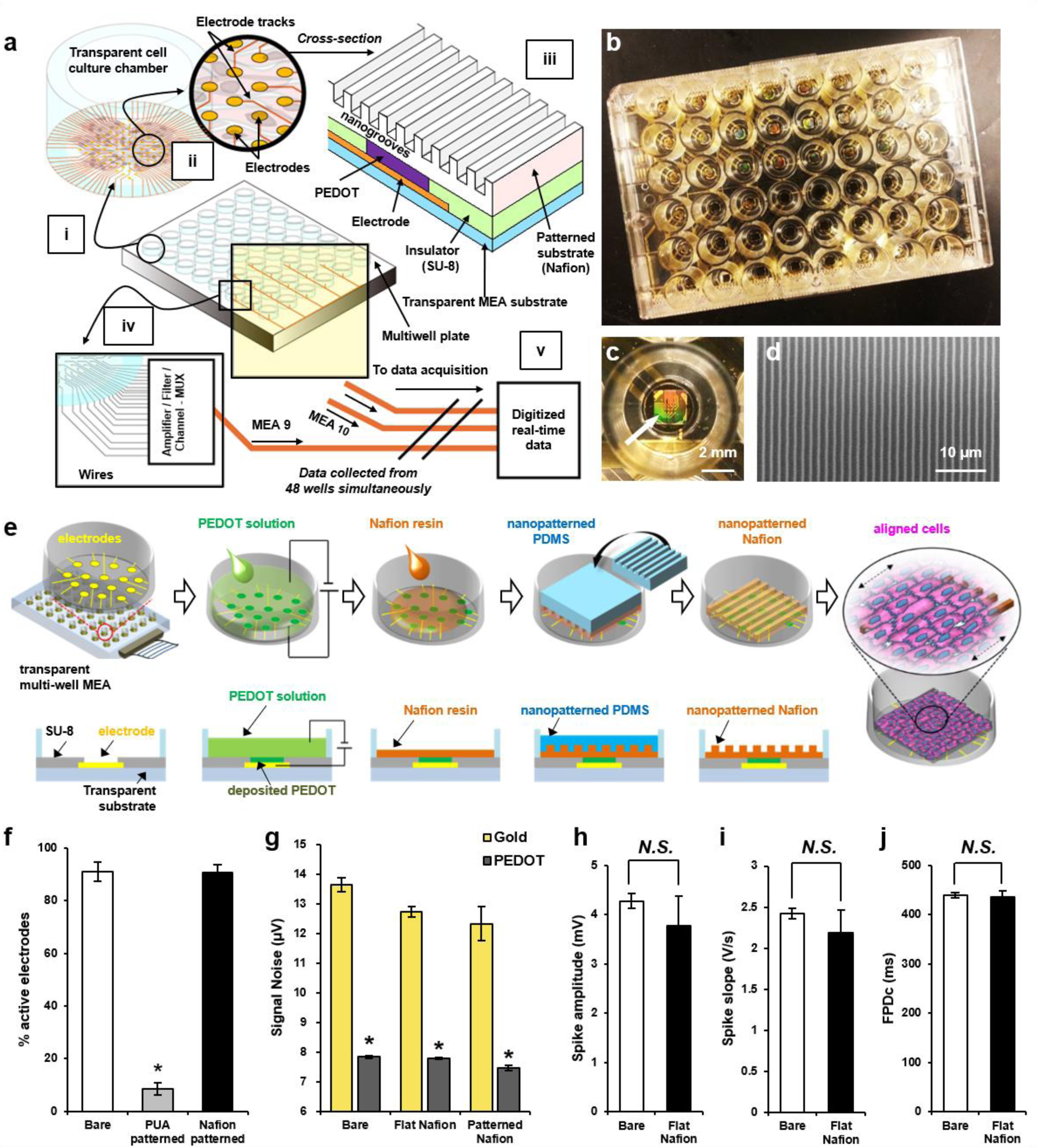
**Design, fabrication, and characterization of the nanotopographically-patterned MEA devices.** (a) High-throughput nanoMEA concept. Each well of the multiwell plate (i) supports independent cell cultures for high-throughput analysis. Within each well (ii), the electrode bed facilitates recording of field potentials generated by the overlying cells. Nanotopography (iii) is applied to each well (or a subset of wells) to promote cellular alignment and functional development. Captured signals are relayed, via an amplifier (iv), to a software program for subsequent analysis (v). (b) Low magnification image of multiwell MEA plate with nanotopography applied to each well. (c) Nafion nanotopography applied to a single well of a 48-well MEA plate. The presence of the nanoscale features causes light diffraction on the surface, giving the patterns a green-orange color in this image (white arrow). (d) SEM image of Nafion nanotopography. (e) NanoMEA fabrication schematic. Pristine wells are first treated with PEDOT to improve the sensitivity of the base electrode. A drop of Nafion resin is then applied to the substrate, and a PDMS mold is pressed into it. After overnight curing, the PDMS mold is removed to reveal Nafion topographic substrates underneath. (f) Percentage of MEA electrodes from which hPSC-CM signal detection could be clearly distinguished above background noise. (g) Quantification of noise recorded from bare, flat Nafion, and patterned Nafion electrodes with and without PEDOT treatment. Cells maintained on untreated and flat Nafion coated MEAs for 7 days were assessed for differences in: (h) depolarizing spike amplitude, (i) depolarizing spike slope, and (j) field potential duration corrected for beat period (FPDc). In all presented data, *p < 0.05.

We used capilary force lithography to create nanotopographic patterns using a polydimethylsiloxane (PDMS) mold cast from a polyurethane acrylate (PUA) master (**Figure 1e**). Preliminary MEA studies using differentiated hPSC-derived cardiomyocyte monolayers on PUA nanotopographic patterns demonstrated that patterned substrates fabricated using materials optimized in previous studies^17, 18^ significantly reduce signal acquisition from underlying electrodes, likely due to the insulating properties of the polymer (**Figure 1f**). To address this issue, the ion-permeable, nanoporous polymer, Nafion, was used to fabricate substrate topographies for subsequent experiments. Given its ionic properties and high thermal and mechanical stability^20, 21^, Nafion enables reliable signal capture of cellullar field potentials from underlying electrodes, while retaining the rigidity required to form high-fidelity nanoscale 3D topographic structures. Additionally, cured Nafion substrates are optically transparent, faciltiating optical assessment of cell cultures prior to funcitonal analysis and *post hoc* immunocytochemical study of cytoskeletal architecture once MEA assessment is complete.

To evaluate whether application of Nafion substrates to MEAs had an impact on signal acquisition, we analyzed impedance measurements and electrode noise on uncoated MEAs, MEAs coated with a flat Nafion layer, and MEAs coated with nanotopographically-patterned Nafion. In addition, recordings were analyzed from electrodes that had been treated with the conductive polymer, Poly(3,4-ethylenedioxythiophene) (PEDOT), prior to Nafion application and those that had not (**Supplementary Figure 1**). PEDOT is used routinely to increase the sensitivity of electrodes^22, 23^ and was investigated here as a means to improve signal capture from the microelectrodes (**Figure 1g**). In all conditions examined, PEDOT application was found to significantly reduce electrode noise and impedance readings, but no difference between uncoated and Nafion coated electrodes was observed. Without PEDOT, electrode noise measurements across 32 independent electrodes were recorded as 13.64 μV ± 0.23, 12.73 μV ± 0.17, and 12.33 μV ± 0.58 for uncoated, flat Nafion, and nanotopographically-patterned Nafion electrodes, respectively. The presence of PEDOT reduced these readings to 7.84 μV ± 0.04, 7.79 μV ± 0.04, and 7.45 μV ± 0.05, respectively. Similarly, impedance magnitude measurements for uncoated (bare), flat Nafion, and nanotopographically-patterned Nafion electrodes went from 2.56 ± 0.58, 1.57 ± 0.16, and 2.41 ± 0.20 MΩ, respectively, to 0.013 ± 0.00027, 0.010 ± 0.00054, and 0.015 ± 0.0019 MΩ following PEDOT addition (**Supplementary Table 1**); a decrease in impedance of over 100-fold.

To evaluate the impact of Nafion coating on signal detection further, electrophysiological recordings were collected from hPSC-CM monolayers after 7 days in culture using uncoated and flat Nafion coated electrodes (**Figure 1h-j**). Nanotopography was not investigated during this validation study as topography-induced alterations in cardiomyocyte electrophysiological properties would confound comparisons between Nafion coated and uncoated substrates. No significant differences in baseline electrophysiological metrics were detected between cells maintained on bare MEAs and those coated with Nafion. As such, the presence of a Nafion layer between the cells and underlying electrodes was determined to have a negligible impact on both the electrophysiological function of cultured cells and the signal capture capabilities of the nanoMEA device.

### Nanotopographically-patterned Nafion substrates promote structural and functional maturation of stem cell-derived cardiomyocytes

To demonstrate that nanotopographically-patterned Nafion substrates were capable of promoting structural maturation of hPSC-CM monolayers to a similar degree as seen previously with single isolated hPSC-CMs^18^ and neonatal rat ventricular myocyte monolayers^17^, immunocytochemical analysis was used to assess cell morphology and sarcomere structure in unpatterned and patterned hPSC-CM cultures (**Figure 2a,b**). Cells on flat substrates were found to exhibit random orientations with poorly organized sarcomeres, whereas nanotopographically-patterned cells displayed anisotropic morphologies with ordered myofibrils and regular z-band alignment. Both sarcomere lengths and z-band widths were found to increase on nanotopographically-patterned substrates (**Figure 2c,d**); an accepted trait for cardiomyocyte maturation^6^. Specifically, sarcomere length in cells maintained on flat surfaces was found to average 1.78 μm ± 0.02 (n = 99), which increased to 1.96 μm ± 0.02 (n = 100) in nanotopographically-patterned cells. Similarly, z-band widths in cultured cardiomyocytes were measured at 5.14 μm ± 0.21 (n = 110) and 6.44 μm ± 0.39 (n = 100) on flat and patterned surfaces, respectively. In addition, quantification of F-actin alignment in unpatterned and patterned cells clearly illustrated the polar nature and anisotropic cytoskeletal organization of cardiomyocytes maintained on nanotopographically-patterned substrates (**Figure 2e**).

**Figure 2:**
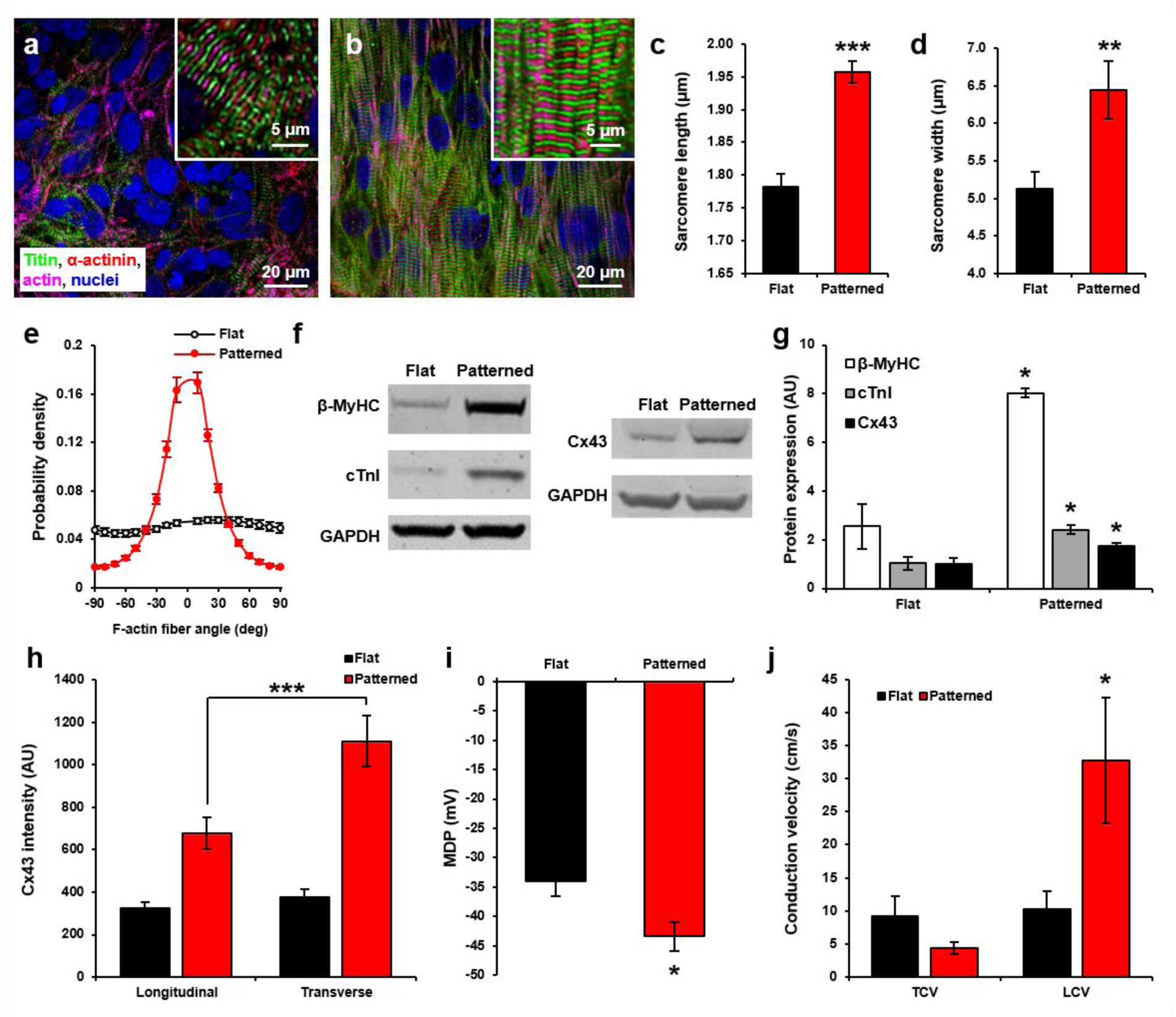
**Enhanced structural and functional properties of human cardiomyocytes on nanoMEAs.** (a) Immunostained image of hPSC-CMs on flat Nafion substrates. Inset shows detail of the sarcomeric structures present. (b) Immunostained image of hPSC-CMs on nanotopographically-patterned Nafion substrates. Inset shows detail of the sarcomeric structures present. (c) Quantification of sarcomere length from patterned and unpatterned hPSC-CMs. (d) Quantification of z-band widths from patterned and unpatterned hPSC-CMs. (e) Histogram detailing frequency of F-actin fibers in cultured cardiomyocytes that possess a given angle of alignment relative to vertical. All images collected from patterned cultures were oriented so that the underlying pattern ran vertically. (f) Immunoblot results from unpatterned and patterned hPSC-CMs, detailing expression levels of β-myosin heavy chain (β-MyHC), cardiac troponin I (cTnI), and connexin 43 (Cx43), as well as GAPDH internal controls. (g) Densitometric analysis of band intensity, providing quantification of the changes in protein expression illustrated in (f). (h) Quantification of pixel intensity in images collected from hPSC-CM cultures stained with a primary antibody that targets Cx43. (i) Measurement of maximum diastolic potential (MDP) using whole cell patch clamp analysis of patterned and unpatterned hPSC-CMs. (j) Measurement of conduction velocity (CV) across hPSC-CM monolayers on flat and nanotopographically-patterned MEAs. Conduction was measured specifically in both the transverse (TCV) and longitudinal (LCV) orientations and underlying nanotopography was organized to run longitudinally. In all presented data, *p < 0.03, **p < 0.003, ***p < 0.001.

Immunoblot analysis of protein lysates collected from unpatterned and patterned hPSC-CMs highlighted that nanotopographic substrates promoted an upregulation of β-myosin heavy chain (β-MyHC), cardiac troponin I (cTnI), and connexin 43 (Cx43) (**Figure 2f,g**). However, no significant difference was observed in levels of slow skeletal troponin I (ssTnI) expressed in cells maintained on flat versus patterned substrates (**Supplementary Figure 2**).

Peroxisome proliferator-activated receptor gamma coactivator 1-α (PGC1-α) expression was found to be significantly increased in cardiomyocytes maintained on nanotopographically-patterned surfaces (**Supplementary Figure 3**). PGC1-α is known to perform a central role in global oxidative metabolism through induction of mitochondrial biogenesis and tuning of the intrinsic metabolic properties of mitochondria^24, 25^. In addition, PGC1-α has also been shown to act as a key regulator of reactive oxygen species (ROS) removal via the expression of several ROS-detoxifying enzymes, including glutathione peroxidase 1 (GPx1) and superoxide dismutase 2 (SOD2)^26^. We therefore performed a fluorometric cellular ROS detection assay on unpatterned and patterned hPSC-CMs to determine whether upregulation of the PGC1-α protein in patterned cells had any notable impact on oxidative stress (**Supplementary Figure 4**). Quantification of superoxide and oxidative stress stains showed no difference between unpatterned and patterned samples, although a trend towards lower levels in patterned cultures was observed for the latter. Patterned cardiomyocytes did exhibit a significant reduction in nitric oxide expression, compared with unpatterned controls, providing support for the hypothesis that nanotopography enhances the oxidative metabolic capacity of hPSC-CMs. In addition, the collected data demonstrate that nanotopographically-patterned culture surfaces exert little effect on the stress state of cardiomyocytes maintained on these surfaces. This point is critical in ensuring that any differences observed in electrophysiological function between unpatterned and patterned cells are due to the maturation state of the cells and not a by-product of a stress response in patterned cultures.

In addition to immunoblot data revealing an upregulation in Cx43 expression in patterned cells, analysis of Cx43 localization in immunostained cultures revealed that Cx43 was more highly expressed in the transverse orientation (relative to the underlying nanotopographic patterns), suggesting that gap junction accumulation was occurring at the polar ends of patterned cells (**Figure 2h; Supplementary Figure 5**). No directional preference for Cx43 expression was observed in unpatterned cells, further highlighting the impact of nanotopography on subcellular organization in hPSC-CMs.

Overall, the anisotropic nature of the cultured cells, the improvements in sarcomere length, myofibril alignment, and gap junction protein expression and localization, as well as the significant upregulation of cardiac specific contractile machinery and metabolic regulator proteins, all serve to underscore the ability for Nafion nanotopographic patterns to enhance the phenotypic development of cultured hPSC-CMs. These results support and build upon those published previously using alternative polymeric materials^18, 27^ and confirm the ability for Nafion nanotopographic patterns to promote the establishment of more physiologically relevant cardiac cell sheets for use in downstream applications.

We then investigated whether the observed changes in hPSC-CM phenotype correlated with any changes in electrophysiological function in these cells. Prior to investigation by MEA, whole cell patch clamp analysis was used to measure membrane potential in single hPSC-CMs. The collected data indicated that patterned cells developed a more negative maximum diastolic potential (MDP) than flat controls (**Figure 2i**), suggesting that patterned cells are able to maintain a more negative resting membrane potential than their unpatterned counterparts; a hallmark of increased cardiomyocyte maturity^6, 28^.

Analysis of MEA field potential recordings revealed that spontaneous baseline beat rate did not differ significantly between unpatterned and patterned hPSC-CM monolayers following 21 days *in vitro.* Critically, analysis of beat-to-beat-variance (through comparison of the median and mean difference in beat interval period from one beat to the next^29^ (**Supplementary Figure 6**) highlighted that both unpatterned and patterned cells exhibited regular and consistent beat intervals at baseline, thereby confirming the presence of stable, non-arrhythmic populations for subsequent assessment. Interrogation of monolayer culture by MEA facilitates assessment of conduction velocity in a manner not possible when using single cell analysis methods, such as patch clamp electrophysiology. For all propagation experiments discussed in this study, longitudinal conduction dictates the vertical orientation on the MEA as all nanopatterns were oriented to run vertically during pattern generation. Conduction velocities measured from hPSC-CMs on our nanoMEA devices were found to be highly anisotropic (**Figure 2j; Supplementary Videos 1 and 2**), thereby more closely mirroring the directed propagation patterns present in the native myocardium. Furthermore, longitudinal conduction velocities measured from patterned cells were substantially faster than similar measurements taken from flat controls, as well as global (non-directional) conduction velocity measurements calculated by the Axion software. Specifically, longitudinal conduction velocity measured from patterned cells was recorded as 32.73 cm/s ± 9.51, whereas flat controls produced longitudinal and global conduction speeds of 10.32 cm/s ± 2.62 and 23.49 ± 4.57, respectively.

### NanoMEAs increase cardiomyocyte sensitivity to cardiotoxic compounds that target structural features of the cell

We sought to establish whether observed differences in cardiomyocyte structural and functional phenotypes led to alterations in cellular responses to cardiotoxic agent exposure. Specifically, we analyzed whether unpatterned and nanotopographically-patterned cells exhibited different sensitivities to compounds with arrhythmogenic or conduction blocking activity. The ability to pattern individual wells within a 48-well MEA plate enabled assessment of multiple doses across both surface conditions within a single device, thereby increasing throughput for such comparative studies. Verapamil was first examined as a negative control compound. As has been reported previously^30, 31^, exposure of unpatterned and patterned cells to increasing doses of verapamil produced a significant shortening of the field potential duration (corrected for beat rate; FPDc) but no notable arrhythmogenic effect (**Supplementary Figure 7**). No significant difference was observed in the response of unpatterned and patterned cells, indicating that the presence of the nanotopography did not induce unexpected arrhythmogenic behavior in cells exposed to this “safe” compound.

Bepridil is a Ca^2+^ release antagonist originally developed to treat angina that has been shown to prolong QT interval in the majority of patients for whom it has been prescribed^32^. The mechanism by which bepridil prolongs QT is yet to be fully elucidated. However, in addition to regulating Ca^2+^ release, it is known that bepridil competes with cTnI for troponin C binding sites, thereby altering Ca^2+^ sensitivity in exposed cells^33^. This suggests that the ability for bepridil to alter cardiomyocyte functional performance may be tied to the presence, organization, and/or function of troponin in these cells. Critically for drug development applications, previous studies with bepridil using conventional MEAs have demonstrated that cultured hPSC-CMs exhibit little electrophysiological response to the compound, even at supra-physiological doses^30^. Since cTnI was found to be upregulated in patterned hPSC-CM monolayers, and the organization of contractile sarcomeres greatly improved in these cells, we investigated whether exposure to bepridil had a more pronounced effect on patterned hPSC-CM monolayers than had been previously reported with conventional MEAs. In this study, treatment of cardiomyocytes maintained on our devices with bepridil led to a significant prolongation of FPDc in patterned cultures, whereas the change in FPDc measured in bepridil-treated flat controls was not significantly different from baseline (**Figure 3a-c**). 1 µM bepridil treatment on unpatterned cells promoted a change in FPDc from 404.46 ms ± 26.44 at baseline to 434.61 ms ± 2.14 (a change of just 7%), suggesting the drug had little impact on the electrophysiology of cardiomyocytes maintained on conventional culture surfaces. Conversely, bepridil treatment on patterned cells was found to increase FPDc from 388.58 ms ± 13.70 at baseline to 463.72 ms ± 2.56 at 1 µM bepridil; an increase of roughly 20%.

**Figure 3:**
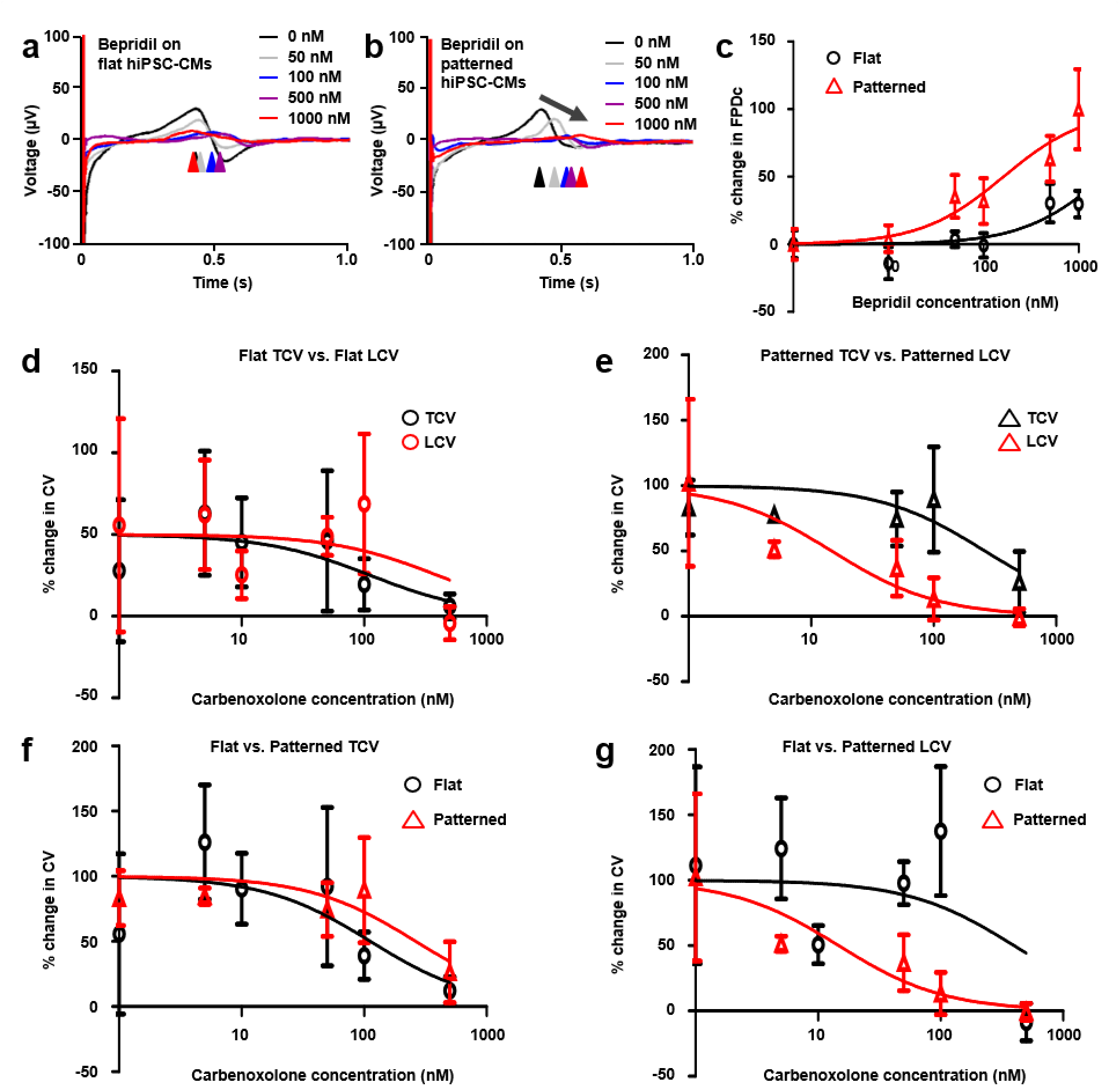
**Electrophysiological response of patterned and unpatterned hPSC-CMs to treatment with bepridil and carbenoxolone.** (a) Representative traces (averaged across 10 beats) recorded from hPSC-CM monolayers on flat MEAs and subjected to increasing doses of bepridil. (b) Representative traces recorded from hPSC-CM monolayers on nanoMEAs and subjected to increasing doses of bepridil. (c) Normalized dose response curve illustrating effect of increasing concentrations of bepridil on the FPDc of unpatterned and patterned hPSC-CMs. The R^2^ values for the unpatterned and patterned cultures were 0.26 and 0.39, respectively. (d) Normalized dose response curves illustrating the effect of increasing concentrations of carbenoxolone on the conduction velocity of unpatterned hPSC-CM monolayers. Dose response curves were calculated from analysis of propagation speeds in both the transverse (TCV) and longitudinal (LCV) directions. The R^2^ values for TCV and LCV curve fits were 0.23 and 0.11, respectively. (e) Normalized dose response curves illustrating the effect of increasing concentrations of carbenoxolone on the TCV and LCV of patterned hPSC-CM monolayers. The R^2^ values for TCV and LCV curve fits were 0.21 and 0.28, respectively. (f) Normalized dose response curves illustrating the effect of increasing concentrations of carbenoxolone on the TCV of patterned and unpatterned hPSC-CMs. (g) Normalized dose response curves illustrating the effect of increasing concentrations of carbenoxolone on the LCV of patterned and unpatterned hPSC-CM monolayers.

Carbenoxolone is a gap junction blocker (currently unavailable in the US) that is typically used to treat ulceration of the gastrointestinal tract. In cardiac tissue, it is known to exert a strong conduction blocking effect, inhibiting action potential propagation and in turn affecting synchronous contraction of the tissue^34^. Analysis of cardiac conduction patterns on our nanoMEA platform highlighted that treatment of cells with carbenoxolone significantly reduced conduction speeds *in vitro* (**Supplementary Videos 3 and 4**). However, substantial differences were observed in the inhibitory effect of the compound, depending on the substrate condition and the orientation of propagation examined. On flat surfaces, no significant difference was observed in the dose response curves generated for transverse conduction velocity (TCV) and longitudinal conduction velocity (LCV). Furthermore, the inconsistency of the propagation waves on these surfaces led to substantial variability in measured values across experimental repeats (n = 4 for each dose examined), making it difficult to construct well-fitted dose response curves (**Figure 3d**). In comparison, TCV and LCV recorded from nanotopographically-patterned cells exhibited significant differences in response to carbenoxolone treatment, with longitudinal propagation affected at significantly lower doses than transverse propagation (**Figure 3e**). Additionally, the consistency of the propagation waves in these cultures created more reproducible data, tighter error bars, and better fitted dose response curves. The standard deviation of the residuals for the fitted data were 55.25 and 74.19 for flat TCV and LCV, respectively, whereas TCV and LCV curves generated for the patterned data had standard deviation of the residuals of 42.10 and 25.15, respectively. Residual values indicate the vertical distance (in Y units; in this case % change in conduction velocity) of the measured points from the fit line. As such, the reduction in standard deviation of the residuals for the patterned data (compared with flat controls) suggests the data points fall closer to the fit line and indicate an overall improvement in the fit. Comparison of TCV values from flat and patterned data sets revealed no significant difference in carbenoxolone dose response (**Figure 3f**). However, analysis of LCV curves from both groups showed increased sensitivity in patterned cells versus flat controls (**Figure 3g**). Specifically, the carbenoxolone IC_50_s, obtained from flat and nanotopographically-patterned LCV data, were calculated as 398.1 nM and 14.71 nM, respectively.

### NanoMEAs increase network connectivity and sensitivity to synaptic blockers in cultured hPSC-neuron monolayers

Given the observed capacity for nanotopographically-patterned MEAs to enhance the functional performance of human cardiomyocytes, we next investigated whether similar improvements in other electrically-active cell types could be achieved via interaction with our Nafion substrates. Human excitatory (glutamatergic) neurons derived from human pluripotent stem cells were maintained on nanoMEAs for 3 weeks before being investigated for changes in spontaneous electrical activity. Image analysis from low density neuronal cultures on flat and nanotopographically-patterned Nafion coated coverslips indicated that, like cardiomyocytes, hPSC-neurons were capable of responding to the anisotropic substrate cues and developed organized neuritic processes that extended from cells in parallel with the underlying topography (**Figure 4a,b**). High density cultures maintained on MEAs exhibited no significant differences in overall firing rates or burst fire behavior across the 3-week culture period examined (data not shown). However, significant changes in network burst behavior between patterned and unpatterned hPSC-neuron populations were detected after 21 days *in vitro* (**Figure 4c-e**). Specifically, spikes per network burst increased from 343.88 ± 33.79 in flat cultures to 496.00 ± 32.49 in cells maintained on patterned surfaces. Spikes per burst per channel likewise increased from 55.95 ± 4.46 on flat surfaces to 74.25 ± 4.26 on patterned substrates. No significant difference was observed in the number of electrodes participating in networks bursts (5.94 ± 0.33 for flat cultures and 6.50 ± 0.35 for patterned cells), indicating that the increase in firing was due to greater activity over each electrode rather than an increase in the number of electrodes participating in the bursts. The average length of network bursts also increased from 1.31 ± 0.05 seconds in flat cultures to 1.52 ± 0.04 in patterned hPSC-neurons, providing further evidence for an increase in synaptic crosstalk in patterned neuronal populations.

**Figure 4:**
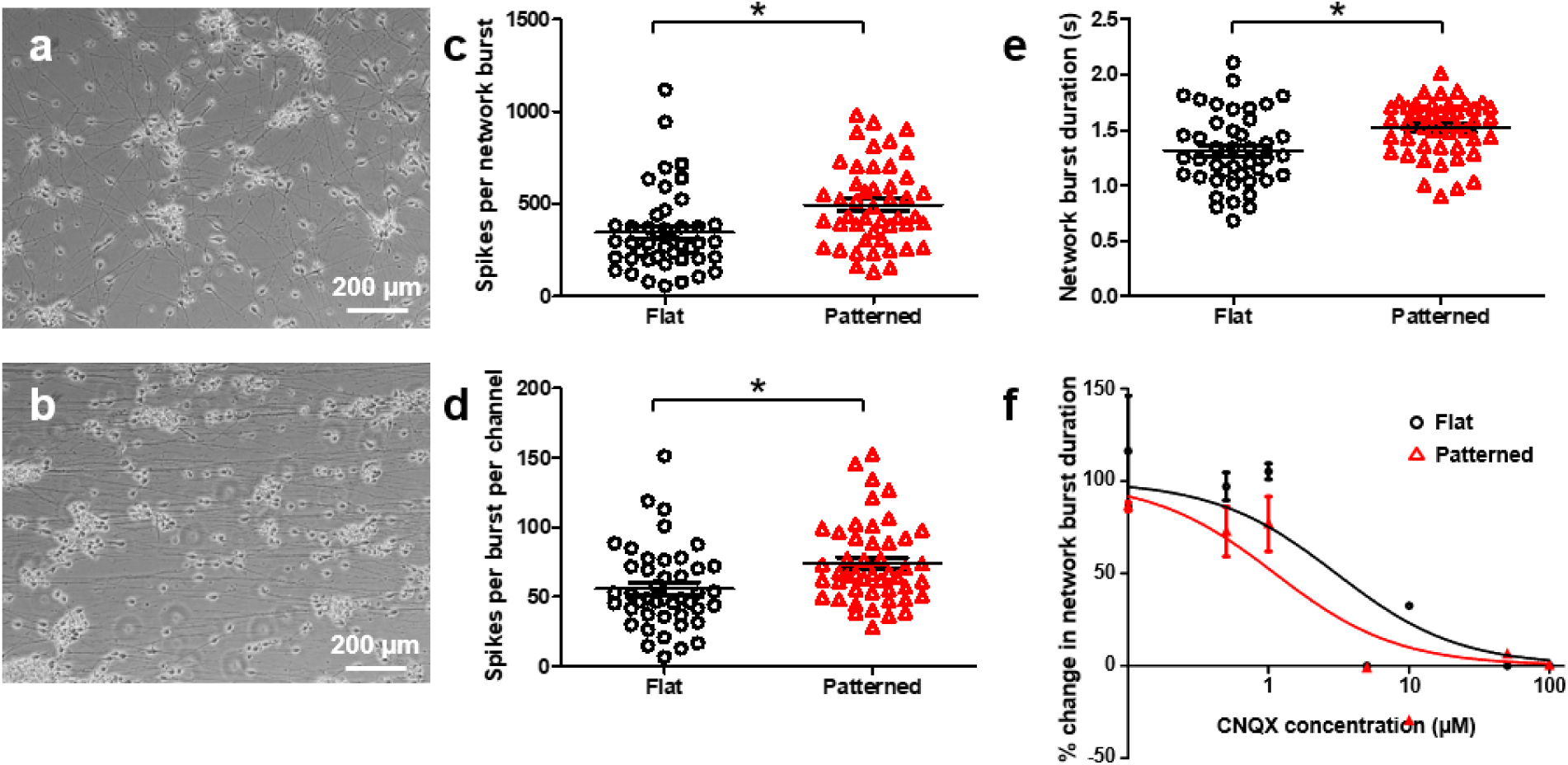
**Electrophysiological responses of hPSC-neurons to culture on nanoMEAs.** (a) Bright-field image of low-density hPSC-neurons maintained on flat Nafion substrates. (b) Bright-field image of low-density hPSC-neurons maintained on nanotopographically-patterned Nafion substrates. (c) Measurement of the number of spikes recorded per network burst from high-density hPSC-neuron populations maintained on flat and nanotopographically-patterned MEAs. (d) Measurement of the mean number of spikes recorded per electrode during individual network bursts from hPSC-neuron populations maintained on flat and nanotopographically-patterned MEAs. (e) Measurement of network burst duration from hPSC-neuron populations maintained on flat and nanotopographically-patterned MEAs. (f) Normalized dose response curve illustrating effect of increasing concentrations of the AMPA receptor blocker CNQX on duration of network bursts in unpatterned and patterned hPSC-neuron populations. The R^2^ values for the unpatterned and patterned cultures were 0.75 and 0.76, respectively. In all presented data, *p < 0.01.

Based on the evidence for increased synaptic activity observed in patterned cells, we sought to establish whether such cultures exhibited altered sensitivity to treatment with synaptic inhibitors. As expected, increasing doses of the AMPA receptor blocker cyanquixaline (CNQX) were found to reduce the duration of network bursts in both substrate conditions (**Figure 4f**). However, the calculated dose response curves suggest that nanotopographically-patterned cells exhibited greater sensitivity to the compound than did their unpatterned counterparts. Statistical comparison of the fitted lines created by nonlinear regression of the normalized data determined that the data was better represented using a separate curve for each data set (p = 0.033), indicating that the observed difference in the dose response curves was significant.

Finally, we looked for additional evidence of structural organization and increased synapse development in patterned hPSC-neuron cultures to support the functional observations described above. Patterned and unpatterned hPSC-neurons grown at low density on coverslips were stained using antibodies against microtubule associated 2 (MAP-2) and neurofilament to observe the organization of dendrites and axons, respectively (**Figure 5a,b**). Interestingly, analysis of the patterned images indicated that while neurofilament positive axons followed the orientation of the underlying topography, MAP-2 positive dendrites did not. Instead, these neurites formed a network pattern across the culture surface with many processes extending in a transverse orientation to the topography. Unpatterned controls showed no organization for either MAP-2 or neurofilament positive neurites, with both neurite types extending randomly across the culture surface. High density hPSC-neuron cultures maintained on flat and patterned surfaces were then stained for expression of pre- (synaptotagmin) and post-synaptic (PSD-95) markers in order to assess levels of co-localization of these proteins (**Figure 5c,d**). Intensity correlation scatter plots were constructed in ImageJ that compared the intensity of each pixel in the red and green channels across entire images collected from these stained cultures. Using this information, we calculated the mean Pearson correlation coefficient (PCC) for each image analyzed as a measure of synaptotagmin and PSD-95 colocalization (**Figure 5e**). The PCC provides a measurement of the linear correlation between two variables and has a value between +1 and −1, where 1 is total positive linear correlation, 0 is no linear correlation, and −1 is total negative linear correlation^35^. The significant increase observed in the patterned cells’ PCC, compared with unpatterned controls, indicates greater consistency in individual pixel intensities between the green and red channels for a given image and therefore suggests greater co-localization of pre- and post-synaptic markers in these cultures.

**Figure 5:**
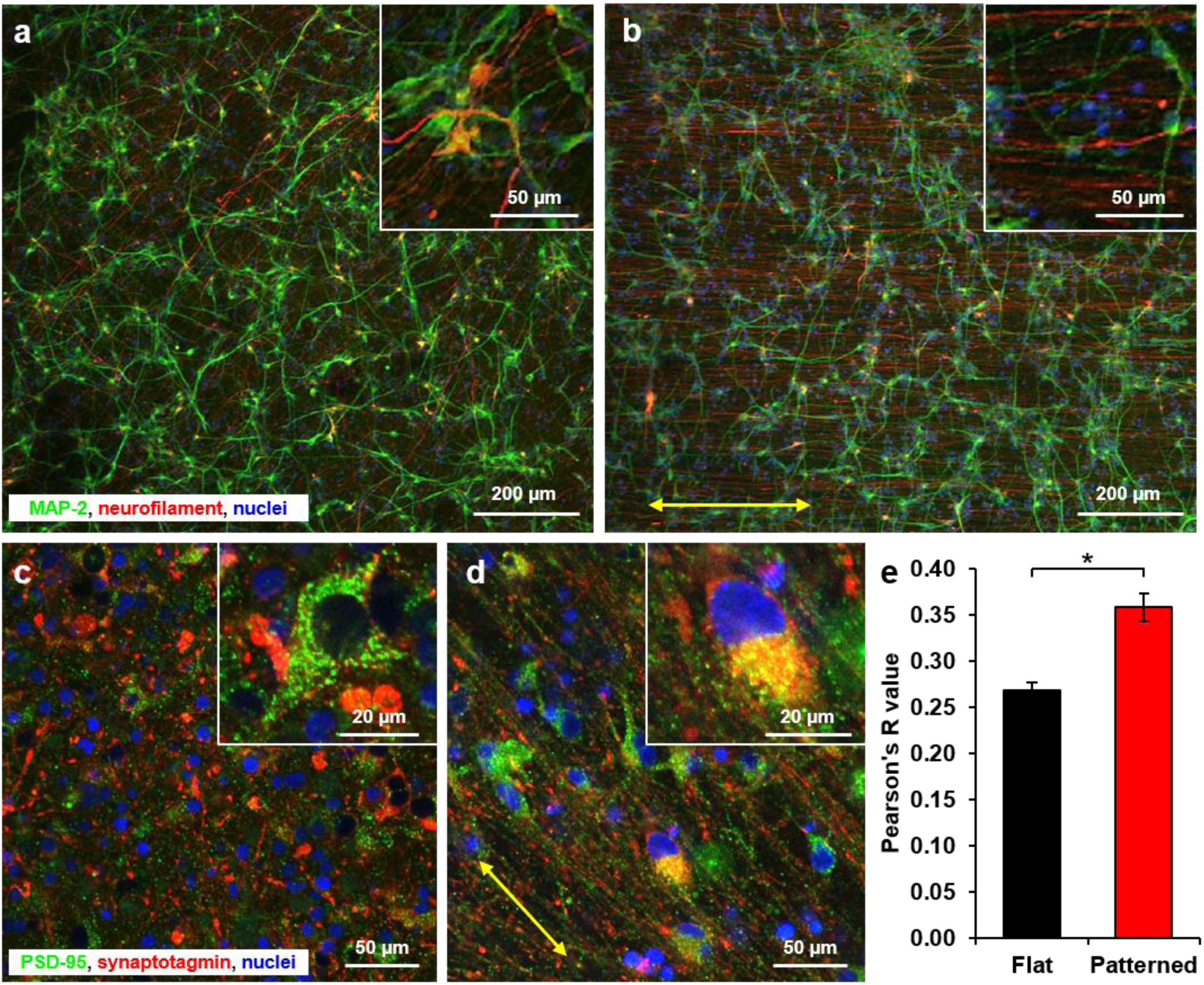
**Structural impact of nanotopographically patterned Nafion substrates on hPSC-neurons.** (a) Human PSC-Ns on flat Nafion substrates immunostained using antibodies against dendrites (MAP-2) and axons (neurofilament) proteins. Inset shows detail of neurite orientation. (b) Human PSC-Ns on nanotopographically-patterned Nafion substrates immunostained using antibodies against dendrites (MAP-2) and axons (neurofilament) proteins. Inset shows detail of neurite orientation. The yellow arrow indicates the orientation of the underlying nanotopography. (c) Human PSC-Ns on flat Nafion substrates immunostained using antibodies against pre- (synaptotagmin) and post-synaptic (PSD-95) proteins. Inset shows detail of the staining associated with a single cell. (d) Human PSC-Ns on nanotopographically-patterned Nafion substrates immunostained using antibodies against pre- (synaptotagmin) and post-synaptic (PSD-95) proteins. Inset shows detail of the staining associated with a single cell. The yellow arrow indicates the orientation of the underlying nanotopography. (e) Pearson correlation coefficients calculated from analysis of scatter plots constructed from images of flat and nanotopographically-patterned hPSC-neurons as an indicator of the relative levels of pre- and post-synaptic marker co-localization. In presented data, *p < 0.002.

## Discussion

Human pluripotent stem cell-derived somatic cells are seen by many as a means to increase the accuracy and downstream utility of preclinical *in vitro* assays. However, the inability of these cells to develop adult-like structural and functional properties has raised questions as to whether such cells are capable of providing preclinical data that is more meaningful than what is currently possible with existing methodologies. The primary objectives of this study were to establish robust, high-fidelity, nanolithographically-patterned microelectrode arrays and to validate these engineered devices in terms of their capacity to enhance the structure and electrophysiological function of hPSC-derived cardiomyocytes and neurons.

Human myocardial tissue possesses a complex structural hierarchy on multiple length scales ranging from macroscopic, tissue-level organization to subcellular nanoscale guidance cues. Based on this understanding of the myocardial structural niche, our group has shown how provision of biomimetic, nanoscale substrate cues, mimicking the size and orientation of myocardial extracellular matrix (ECM) fibers, can be used to promote the structural and functional development of cultured rodent and human cardiomyocytes^17, 18^. In this study, our data demonstrate how ion-permeable nanotopographic patterns can be utilized to attenuate the structural development of hPSC-CM monolayers on MEAs. The use of the ion-permeable polymer Nafion to fabricate the described nanotopographic features constitutes a simple, cost-effective, and reproducible means to organize cardiomyocyte development *in vitro*. The nanoporous nature of the Nafion polymer facilitates efficient signal exchange between underlying electrodes and overlying cells, enabling effective analysis of field potential properties in cultured cardiac monolayers.

Previous analysis of ssTnI and cTnI as a quantifiable ratio-metric maturation signature for induced pluripotent stem cell-derived cardiomyocytes has highlighted that such cells express ssTnI consistently throughout long-term (9-month) culture, whereas expression of cTnI increases over this period^36^. In the current study, the significant upregulation of cTnI protein in cells maintained on nanotopographically-patterned surfaces suggests that substrate topography was able to promote substantial sarcomeric maturation over the 3-week culture period examined. The increase in cTnI expression after 3 weeks in nanotopographically-patterned culture correlates with the increased expression in unpatterned cells after 2, 6, and 9 months *in vitro* reported previously^36^. As such, the troponin protein expression data presented here, coupled with the observed improvements in cellular alignment and sarcomeric development, indicate that nanotopographically-patterned Nafion substrates promote more rapid maturation of hPSC-CM contractile machinery compared with cells maintained on conventional flat surfaces. This observation was further supported by whole cell patch clamp data that demonstrated how patterned Nafion substrates could be used to promote the development of hPSC-CMs with more negative maximum diastolic potentials; a common functional indicator of cardiomyocyte maturity^6^.

Using our 48-well nanoMEA platform, we were able to perform high-throughput analysis of the electrophysiological properties of nanotopographically-patterned hPSC-CM monolayers. The presented data demonstrate that biophysical regulation of cardiomyocyte structural maturation, via presentation of nanoscale topographic cues, leads to concurrent changes in functional output. Specifically, nanotopographically-patterned Nafion substrates promote the development of conduction patterns and speeds in hPSC-CM monolayers that more accurately recapitulate those found in mature human cardiac tissue. The recorded polarization of gap junction proteins in patterned cell monolayers, coupled with cardiomyocyte alignment and elongation in parallel with the underlying topography, likely account for the anisotropic propagation waves observed. Furthermore, they demonstrate how recapitulation of native myocardial architecture can help cultured cardiac monolayers function in a manner more representative of the *in vivo* human tissue.

The calcium channel blocker, bepridil, was found to exert a more powerful FPDc prolongation effect in patterned versus unpatterned cells. The capacity for bepridil to elicit a more pronounced effect on cultured cardiac monolayers may be attributable to its interaction with cytoskeletal elements within cardiomyocytes. Previous work has demonstrated that bepridil accumulates within cardiac cells and binds tightly to F-actin^37^. More recently, it has been shown that bepridil competes for troponin C binding sites with cTnI, which alters Ca^2+^ sensitivity in exposed cells^33^. The fact that bepridil interacts closely with cytoskeletal elements and contractile proteins, which we have shown to be upregulated and reorganized in patterned cardiomyocyte populations, may provide an indication as to why this drug elicited altered responses when applied to cells maintained on nanoMEA devices. If correct, this observation underscores the importance of correctly orchestrating cytoskeletal development and structural maturation within hPSC-CM monolayers to ensure correct physiological responses to treatment with specific classes of drugs.

Analysis of transverse and longitudinal conduction velocity (TCV and LCV, respectively) in patterned hPSC-CM monolayers demonstrated that propagation was significantly more anisotropic in patterned cells. Moreover, comparison of results for carbenoxolone-induced changes in the LCV and TCV of patterned and unpatterned monolayers highlighted that patterned cells exhibit increased, and more consistent, sensitivity to the conduction-blocking compound. The fact that Cx43 was more highly expressed in patterned cells, and localized preferentially at cardiomyocytes’ polar ends, may explain the altered ability to detect carbenoxolone effects when examining propagation speeds in patterned and unpatterned cells and along different orientation planes. The IC_50_ for carbenoxolone calculated from LCV measurements in patterned cells was found to be 14.71 nM; roughly 27-fold lower than results calculated from unpatterned LCV measurements. Carbenoxolone plasma concentrations in patients vary substantially, with numbers in the 55.06 μM ± 21.76 range^38^. However, clinically relevant unbound concentrations are considerably lower; and have been reported in the low nM range^39^. The fact that the carbenoxolone IC_50_ calculated from flat LCV data was measured at 398.1 nM, suggests that flat hPSC-CM monolayers require higher than physiological doses of the drug before an effect can be reliably detected. Critically, our results indicate that structural organization of the cardiac monolayer, enabling study of longitudinal propagation specifically, facilitates detection of compound action at lower doses, thereby offering a more physiologically accurate recapitulation of carbenoxolone activity *in vivo*. Collectively, the bepridil and carbenoxolone data presented in this study suggest that our nanoMEA platform may help improve the predictive capacity of current MEA-based preclinical screening paradigms in terms of their ability to model human myocardial responses to drug exposure.

As was observed in cardiac cultures, hPSC-neurons cultured for 3-weeks on nanoMEAs were found to exhibit improved functional parameters when maintained on patterned surfaces compared with flat controls. Network bursting on MEAs describes incidences where a burst of activity detected on one electrode synchronizes with burst activity detected on other electrodes, and serves as indication of synaptic crosstalk in the cultured cells. The increase in network burst duration and number of spikes per burst potentially indicates an increase in synaptic density in patterned cultures, facilitating longer periods of synaptic activity between groups of neurons following spontaneous activation of burst behavior. Calculated IC_50_ values for the AMPA receptor blocker CNQX in our neuronal experiments dropped from 3.03 µM in flat cultures to 1.12 µM in patterned cells, providing additional evidence for increased synapse presence in nanotopographically-patterned hPSC-neurons. Furthermore, binding affinity experiments previously conducted *in vivo,* have detailed IC_50_s for CNQX in mammalian brains as low as 300 nM^40^. The fact that a lower IC_50_ value was calculated from nanotopographically-patterned hPSC-neurons and that this result was closer to the IC_50_ reported for CNQX *in vivo* highlight the potential for nanotopographically-patterned substrates to enhance the development of neuronal cultures to the point where more accurate predictions of CNS drug effects may be achievable. As a means to explain the observed change in CNQX sensitivity, we examined images of cells stained for expression of pre- (synaptotagmin) and post-synaptic (PSD-95) markers. Analysis of the mean Pearson correlation coefficient (PCC)^35^ for these images highlighted a greater correlation of pixel intensity in images of patterned cells than unpatterned controls, suggesting greater incidence of co-localization of green and red staining. This in turn suggests that greater incidence of pre-and post-synaptic marker co-localization occurred in patterned cultures indicating a possible increase in synapse formation and so greater sensitivity to synaptic antagonists such as CNQX. Staining of low-density coverslips coated with either patterned or unpatterned Nafion indicated that neurofilament positive axons exhibited greater affinity for the topographic cues than MAP-2 positive dendrites. This observation may indicate that while developing axonal growth cones respond to topographic guidance cues, dendritic trees are more heavily influenced by other factors, perhaps the release of factors from neighboring cells. In such a model, the spread of dendritic trees toward neighboring cells would create a network well suited to receiving incoming axons guided by uniaxial topographic cues. Such a system would, to a degree, model the layered structure of the cortex, with axons running uniaxially between cortical layers, and could help explain the ability for patterned cultures to develop denser synaptic connections than randomly organized cells on flat surfaces.

In conclusion, nanotopographically patterned Nafion substrates exert a strong influence on hPSC-derived cardiomyocyte and neuron structural development. Use of our novel nanoMEA device confirmed that these structural changes correlate with altered electrophysiological behavior in these cell populations that may mimic native tissue function more closely. Improved structural development was found to enhance hPSC-CM sensitivity to known cardiotoxic compounds that interact with structural elements within the cell and to alter hPSC-neuron responses to treatment with the synaptic inhibitor CNQX. Based on these findings, the nanoMEA represents an exciting new tool for studying structure-function interplay in excitable human cell populations, and may hold value in improving current preclinical screening assays based on electrophysiological analysis.

## Materials and Methods

### Fabrication of nanotopographically-patterned MEA devices

Traditional multiwell MEAs utilize a PCB-based manufacturing process, which produces an opaque device that prevents optical observations with an inverted microscope. In order to address this shortcoming, we developed a novel 48-well transparent microelectrode array that permits observation of the cells cultured on the arrays. This transparent microelectrode array is composed of a 150-µm thick transparent polymer substrate with gold traces. A thin layer of SU-8 defines 50 µm diameter electrode openings, which are electropolymerized with a conductive polymer, poly(3,4-ethylenedioxythiophene) or PEDOT.

Nanotopographically-patterned Nafion features within this MEA platform were fabricated using capillary force lithography. First, nanogrooved polyurethane acrylate (PUA) master molds were fabricated using a replica molding process, as described previously^17^. Polydimethylsiloxane (PDMS; Sylgard 184, Dow Corning) was then mixed at a 10:1 ratio (base: curing agent). The PDMS solution was applied to the PUA mold and cured overnight at 65°C. Cured PDMS nanotopographic patterns were then removed from the PUA mold, cut to the desired size, and pressed into a 2 µL drop of Nafion resin deposited onto the desired MEA substrate, where capillary action forced the Nafion solution to conform to the PDMS topography. Nafion-coated PDMS patterns were pressed onto MEA surfaces and left for 48 hours to cure via solvent evaporation. All nanotopographic patterns were oriented to run vertically to enable effective comparison between wells. After curing, the PDMS molds were removed, revealing thin Nafion films supporting the desired nanotopography. Prior to use in any cell-based experiments, pattern fidelity of the nanofabricated substrates was assessed by scanning electron microscopy (SEM, SU8000, Hitachi). SEM images were obtained using an accelerating voltage of 5.0 kV following platinum coating of the samples. Impedance measurements were collected prior to cell based experiments using an Omnicron Labs Bode 100 Network Analyzer (Houston, TX), and were collected with reference to the MEA ground.

### Cell culture

Commercially sourced (iCell) hPSC-CMs and hPSC-neurons were purchased from Cellular Dynamics International (CDI) and were stored, thawed, and maintained according to the manufacturer’s protocol. Cultures were maintained in a 37°C/ 5% CO_2_ environment throughout the culture period.

Establishment of optimal cell cultures on nanoMEAs and conventional MEA controls involved drop seeding of high-density cell solutions directly over the desired recording electrodes. A 5 μg/ mL fibronectin coating was first applied for 1 hour to all MEA substrates for cardiomyocyte culture, whereas a 5 μg/ mL laminin coating was used for neuronal cultures. The solution was applied as a 6 μL droplet and positioned to cover just the recording electrodes. Distilled H_2_O was plated around each MEA well to humidify the plate and prevent biopolymer evaporation. Following incubation, the biopolymer was aspirated and either 17,500 cells (cardiomyocytes) or 100,000 cells (neurons), resuspended in 6 μL of medium, were transferred to the biopolymer-coated electrodes. MEAs were then incubated at 37°C, 5% CO_2_ for 3 hours to facilitate cell attachment. At this point, additional medium was added to the MEA well to support the cultured cells thereafter. Cultures were maintained for 21 days prior to analysis (unless otherwise stated), and medium was replaced every 2-3 days during this period.

### Electrophysiological assessment of cardiomyocyte and neuron function

48-well MEA plates were used to assess cardiac and neuronal electrophysiological properties prior to (baseline) and 30 minutes after drug treatment using the Maestro MEA system (Axion Biosystems). During data acquisition, standard recording settings for spontaneous cardiac field potentials or neuronal spikes were used (Axis software, version 2.1), and cells were maintained at 37°C/ 5% CO_2_ throughout the 2-minute recording period. The standard settings have 130 × gain, and record from 1 to 25 000 Hz, with a low-pass digital filter of 2 kHz for noise reduction. For cardiomyocytes, the beat detection threshold was 300 µV, and the field potential duration (FPD) detection threshold was 1.5 × noise. Field potentials were analyzed with the platform software for the following outputs: beat period, fast Na^+^ slope (V/s), fast Na^+^ amplitude (mV), FPD (ms), and conduction velocity (cm/s). For neurons, spike detection was set at 5x the standard deviation of the noise and network burst detection was recorded if at least 10% of the electrodes in a given well showed synchronous activity. Reported results were calculated by averaging all of the electrodes in each well, then averaging data from duplicate wells. Data from drug-treated samples were normalized to baseline initially, then expressed as a percent change from vehicle-treated wells. *Post hoc* analysis of cardiac data was used to create beat map images to illustrate conduction patterns. For this, a custom written MATLAB program was prepared that measured the time delay for each electrode in comparison to the activation point for a given beat. Each electrode was then color-coded accordingly to provide an indication of the propagation pattern. The code also calculated conduction velocity component magnitudes to enable direct comparison of longitudinal and transverse conduction velocities within a single MEA substrate.

In addition to MEA analysis, whole cell patch clamp was performed on individual cardiomyocytes to measure maximum diastolic potential. Whole-cell patch clamp recordings were performed on the 37°C heated stage of an inverted DIC microscope (Nikon) connected to an EPC10 patch clamp amplifier and computer running Patchmaster software (HEKA). Patterned and unpatterned coverslips were loaded onto the stage and bathed in a Tyrode’s solution containing 140 mM NaCl, 5.4 mM KCl, 1.8 mM CaCl_2_, 1 mM MgCl_2_, 10 mM glucose, and 10 mM HEPES. The intracellular recording solution (120 mM L-aspartic acid, 20 mM KCl, 5 mM NaCl, 1 mM MgCl_2_, 3 mM Mg^2+^-ATP, 5 mM EGTA, and 10 mM HEPES) was loaded into borosilicate glass patch pipettes (World Precision Instruments). Patch pipettes with a resistance in the range of 2–6 MΩ were used for all recordings. Offset potentials were nulled before formation of a gigaΩ seal and fast and slow capacitance was compensated for in all recordings. Membrane potentials were corrected by subtraction of a 15 mV tip potential, calculated using the HEKA software. Gap-free recordings of spontaneous activity of patched neurons were performed for 30 seconds with 0 pA current injection to provide a measure of the maximum diastolic potential held by the cell without current input. Cells that required more than 100 pA of current to achieve a −70 mV resting membrane potential were excluded as excessive application of current is indicative of poor patch quality and/or membrane integrity.

### Immunocytochemistry and structural analysis

To verify cellular alignment and structural enhancement on nanotopographically-patterned Nafion, cardiomyocytes were immunostained for markers of cardiac contractile machinery, whereas neurons were stained for evidence of neuritic outgrowth and synaptic density. Briefly, cells were fixed in 4% paraformaldehyde for 15 minutes and blocked with 5% goat serum in PBS for 1 hour at room temperature. Cells were then incubated with primary antibodies diluted in 1% goat serum in PBS overnight at 4°C. The next day, cells were washed 3 times with PBS. They were then incubated in a secondary antibody solution containing secondary antibodies and fluorescently-labelled phalloidin (for F-actin visualization; 1:200, Invitrogen) diluted in 1% goat serum in PBS. Counterstaining was performed with Vectashied containing DAPI (Vector Labs). Images were taken at the Garvey Imaging Core at the University of Washington’s Institute for Stem Cell and Regenerative Medicine using a Nikon A1 Confocal System on a Ti-E inverted microscope platform. A 60x CFI Plan Apo NA1.2 objective lens was used. 12-bit 1024×1024 pixel images were acquired with Nikon NIS Elements 3.1 software. For all images, the pinhole was set to 1 Airy unit. Antibodies used in this study were as follows: mouse anti-α-sarcomeric actinin monoclonal antibody (1:500, Sigma-Aldrich), rabbit anti-M-line titin (M8/M9 epitope) monoclonal antibody (1:300, Myomedix), rabbit anti-connexin 43 monoclonal antibody (1:500, Sigma-Aldrich), mouse anti-neurofilament (1 in 500, Millipore), rabbit anti-MAP-2 (1 in 1000, Millipore), mouse anti-synaptotagmin (1 in 500, Developmental Studies Hybridoma Bank), rabbit anti-PSD-95 (1 in 300, Cell Signaling Technology) Alexafluor-594 conjugated goat-anti-mouse secondary antibody (1:200, Invitrogen), and Alexafluor-488 conjugated goat-anti-rabbit secondary antibody (1:200, Invitrogen).

Sarcomere lengths and z-band widths were measured using standard analysis plugins in ImageJ. First, a straight line was drawn perpendicular to at least ten consecutive well-defined α-actinin-positive bands. An intensity profile was then plotted across the entire length of the line and the length of the band, as well as the distance between intensity peaks, was calculated. For each sample, 100 measurements were collected across 10 random fields of view. Actin alignment and cellular anisotropy was investigated using a custom MATLAB script to assess fluorescently labelled F-actin fibers as described previously^41^. A 2D convolution was performed on each image and a Sobel edge-emphasizing filter was applied to extract the horizontal and vertical edges within the image. These edges were then combined to calculate the gradient magnitude of each pixel. A magnitude threshold was applied to select the major F-actin fibers. Next, the gradient orientation was calculated by determining the angle of the gradient with the x- and y-axes. Finally, the angle that the orthogonal of the gradient makes with the x- and y-axes was calculated. This orthogonal angle (from −90° to +90° with respect to the x-axis) constitutes the principle orientation of the individual F-actin fibers. A probability density histogram of the fiber angle distribution was calculated for each image and averaged per condition. C×43 intensity in transverse and longitudinal orientations was calculated using ImageJ. A 36 × 15 μm region of interest was drawn around the brightest area of staining for each axis for 10 independent cells across multiple images from unpatterned and patterned samples. Average fluorescence intensity for each region of interest drawn was calculated by ImageJ and averaged to give a mean indication of Cx43 intensity in each orientation. In all Cx43 images, transverse measurements were taken horizontally and longitudinal measurements were taken vertically. All images of patterned cells were aligned so that nanotopographic patterns ran vertically. Similarly, brightness of staining for synaptic proteins, as evidence of synaptic density, was performed by constructing intensity correlation scatter plots and calculating the Pearson correlation coefficient for each analyzed image using the Coloc 2 plugin for ImageJ. The flurometric cellular ROS detection assay (AbCam) was carried out using the stains provided in the kit and according to the manufacturer’s protocol.

### Extraction of cellular proteins and immunoblot analysis

Cardiomyocytes were seeded on flat and patterned Nafion substrates at 100,000 to 140,000 cells/ cm^2^ and maintained in culture for 20 to 21 days. Cellular proteins were extracted with M-PER mammalian protein extraction reagent (Thermo Scientific) freshly supplemented with a protease inhibitor cocktail (Sigma-Aldrich, St. Louis, MO) and reducing agent (Invitrogen, Grand Island, NY). The concentration of protein was quantitated using a Pierce BCA Protein Assay Kit (Pierce Biotechnology, Rockford, lL). Immunoblot analysis was performed using a protocol modified from one described previously^42^. Proteins were fractionated in 4-12% NuPAGE Bis-Tris gels (Invitrogen) and transferred to nitrocellulose or PVDF membranes. The primary antibodies used were raised against the following proteins: Connexin-43 (Cx43; C6219, Sigma-Aldrich), inward-rectifier potassium ion channel (Kir2.1; AB85492, Abcam, Cambridge, MA), K_v_ 11.1 (AB136467, Abcam), cardiac β-myosin heavy chain (MyHC; A4.951, Developmental studies hybridoma bank, University of Iowa, Iowa City, IA), cardiac Troponin I (cTnI; AB52862, Abcam), slow skeletal Troponin I (ssTnI; NBP2-46170, Novus Biologicals), peroxisome proliferator-activated receptor gamma coactivator 1-alpha (PGC-1α; AB54481, Abcam), and glyceraldehyde 3-phosphate dehydrogenase (GAPDH; sc-48166, Santa Cruz Biotechnology, Dallas, TX). Membranes were stripped using Reblot plus a mild antibody stripping solution (Millipore, Billerica, MA). The blots were detected with Western Lightning Plus–ECL (Perkin Elmer Life Sciences, Waltham, MA) and photographed using a digitalized FluorChem Q multi image III system (Alpha Innotech/Protein Simple, Santa Clara, CA). The intensity of each protein band was quantified using Photoshop and the values were normalized to the intensity of GAPDH.

### Drug treatments

All drugs were dissolved in dimethylsulphoxide (DMSO) to create a working stock solution. They were then diluted in culture medium so that the highest DMSO concentration exposed to the cells was 0.001%. Drug solutions were made up at 10X the desired concentration. After baseline MEA recording, one tenth of the medium was removed from the cells and replaced with the medium containing the 10X drug compound to bring down the concentration to 1X. This method was employed to reduce drug exposure shock and ensure the cells were still active for subsequent recording. Drug solutions were incubated with cells for 30 minutes before recordings were taken. For each drug, a DMSO only control sample was recorded and all observed changes were expressed as percent changes from these carrier control samples. Each drug was tested at 3 or more concentrations, with a separate well used for each treatment.

### Statistical analyses

Experiments were performed in triplicate, and repeated at least three times using independent vials of cells, with the exception of cardiomyocyte immunoblot experiments, which were collected from two independent experiments. During platform validation, experimental groups consisted of bare, flat Nafion, and nanotopographically-patterned Nafion. For drug sensitivity experiments, each condition was tested on flat Nafion and nanotopographically-patterned Nafion only. Significant differences between groups were evaluated using either unpaired t-tests or Mann Whiney-U tests depending on whether the data passed normality tests. ANOVA, with *post hoc* tests for multiple comparisons, was used to compare data sets with more than two experimental groups. Statistical comparison of dose response curves was conducted using an extra sum of squares F test. Dose response curves were normalized by expressing untreated values as 0% change, and the % change observed in patterned samples at the highest dose tested as 100% change. All other data points were then expressed relative to that scale. In all experiments, a p value of less than 0.05 was considered significant. All statistical tests were performed using the GraphPad Prism statistics software.

## Acknowledgements

This work was supported by NIH R01 HL135143, NIH R01 NS094388, and NIH UG3 EB028094 (D-H. Kim), NIH R24 HL117756, NIH R01 HL126527 and NIH R01 HL130020 (J.C. Wu), and an NIH T32 post-doctoral fellowship T32 HL007312 (A.S.T. Smith).

## Supplementary Figures

**Supplementary Figure 1:**
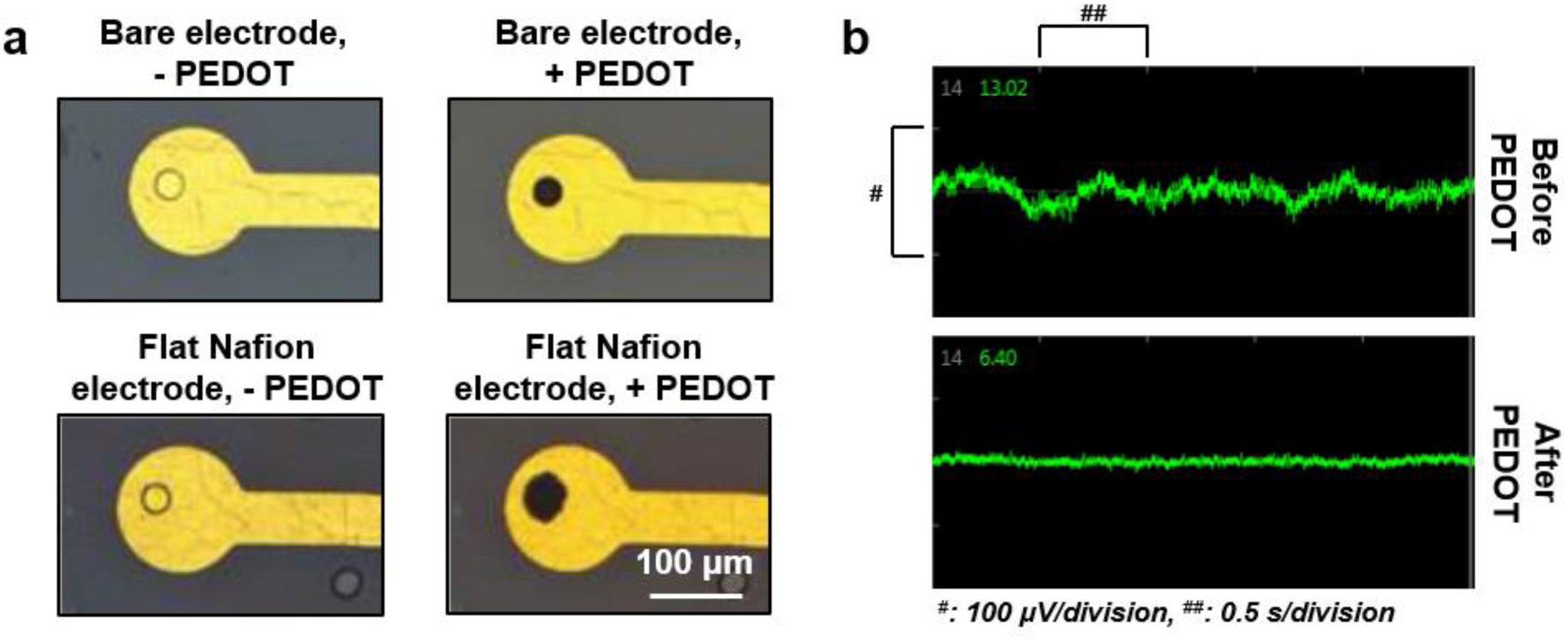
**PEDOT coating on bare and Nafion coated MEAs.** (a) Representative images of bare and Nafion coated electrodes with and without PEDOT treatment. (b) Representative noise recordings from MEAs made in saline solution.

**Supplementary Figure 2:**
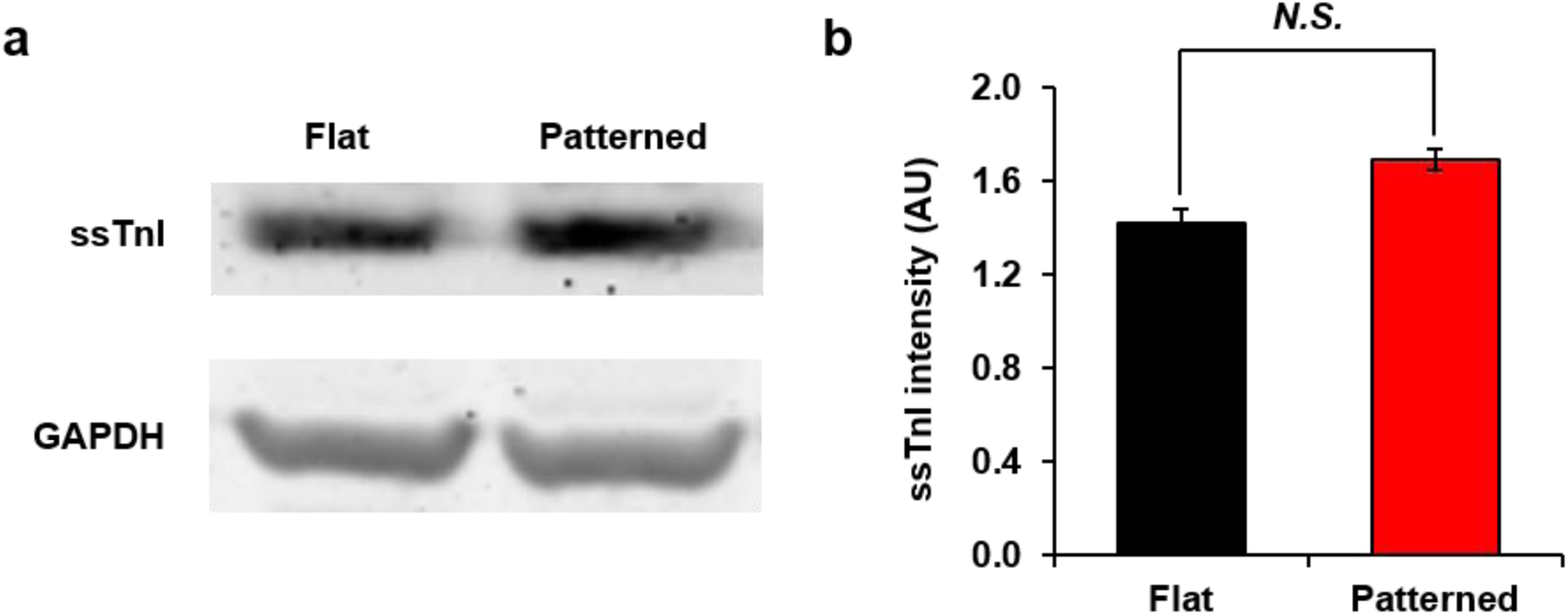
**Slow skeletal troponin I protein expression from unpatterned and patterned hPSC-CM cultures.** (a) Immunoblot blot results illustrating expression of ssTnI, as well as GAPDH internal control, in cardiomyocytes maintained on flat and patterned surfaces. (b) Densitometric analysis of ssTnI western blot results. Data is normalized to GAPDH and expressed in arbitrary units (A.U.). No significant difference was observed between conditions (p = 0.33).

**Supplementary Figure 3:**
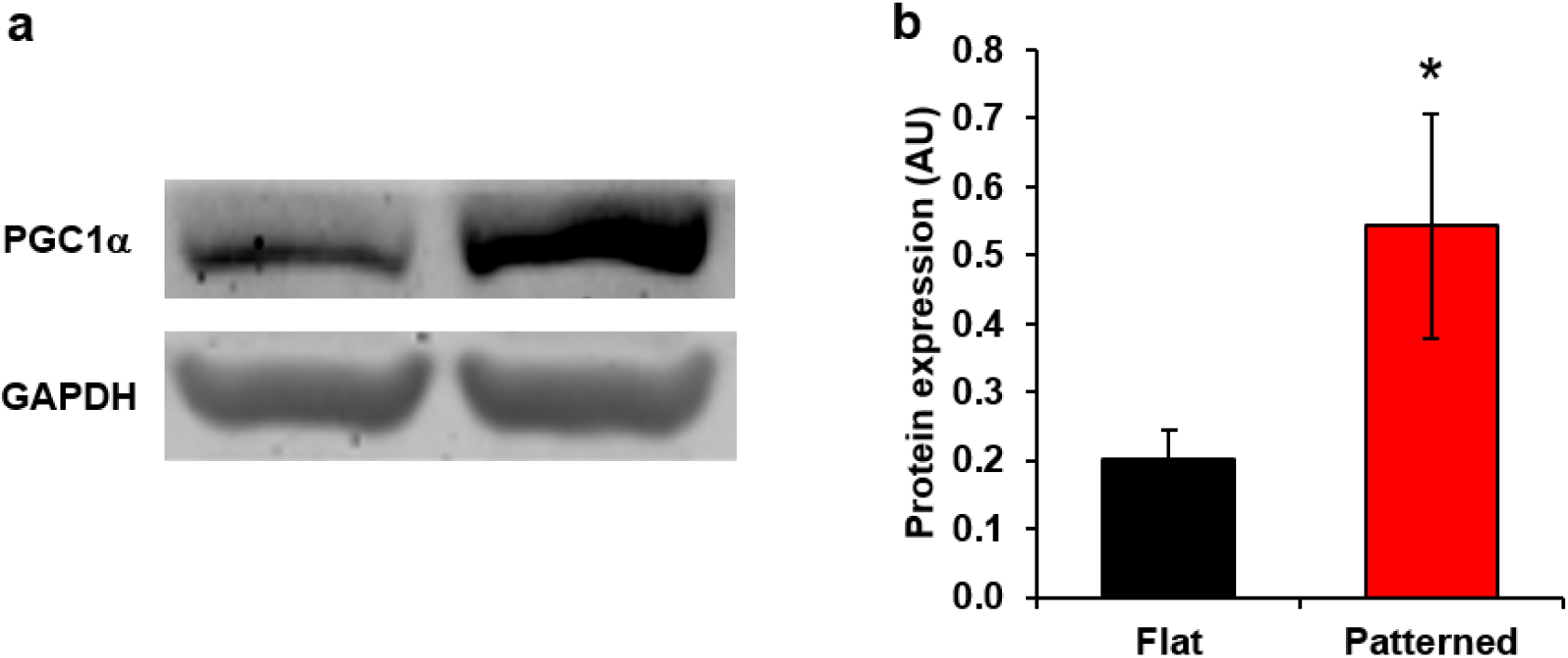
**PGC1α expression in hPSC-CMs maintained on flat and nanotopographically-patterned substrates.** (a) Immunoblot results from unpatterned and patterned hPSC-CMs, detailing expression levels PGC1-α and GAPDH internal controls. (b) Densitometric analysis of band intensity, providing quantification of the changes in protein expression illustrated in (a).

**Supplementary Figure 4:**
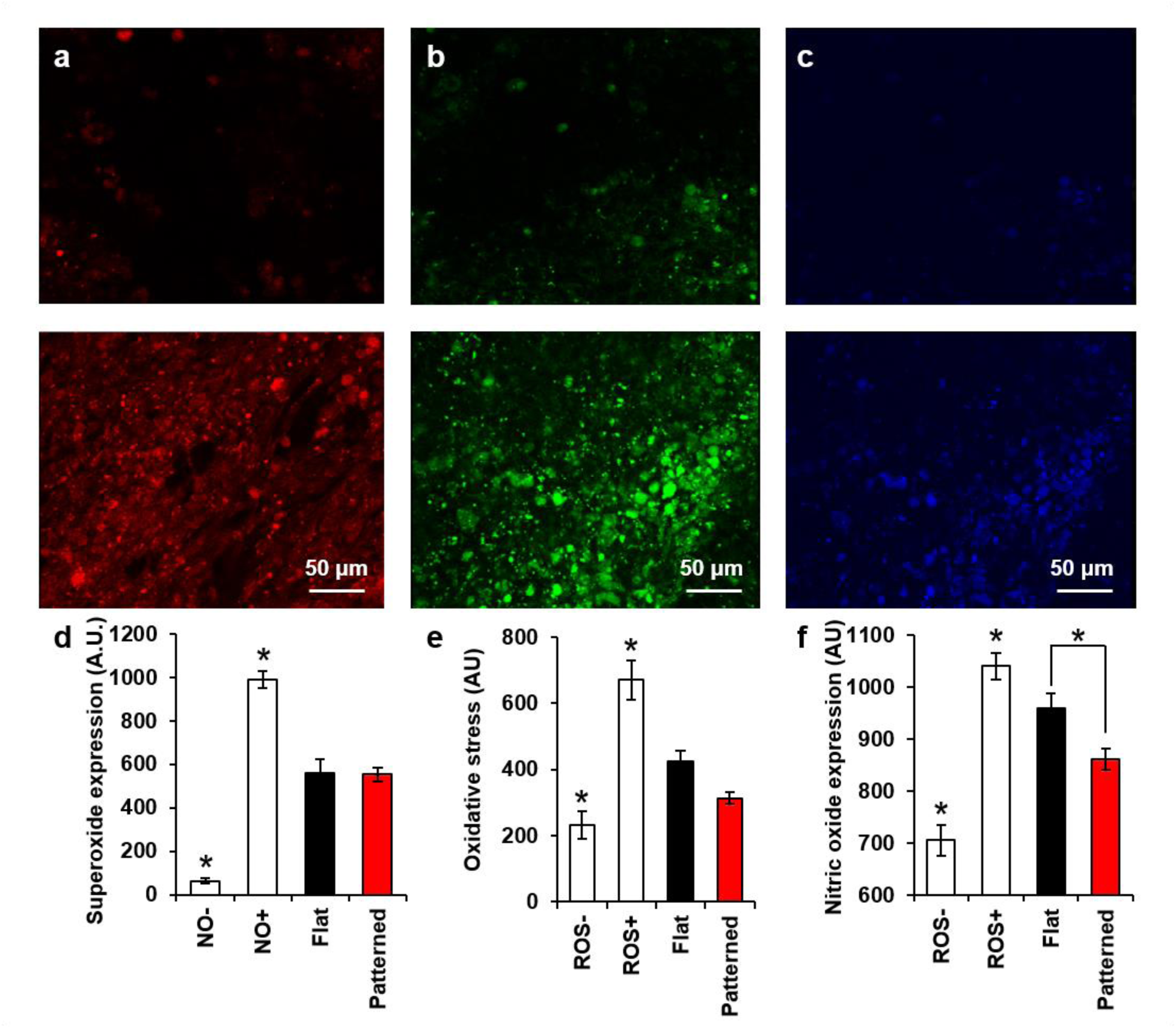
**Effect of nanotopography on reactive oxygen species production in cultured hPSC-CMs.** (a) Representative image from patterned cardiomyocyte culture stained with a superoxide detection reagent (red) following treatment with a reactive oxygen species (ROS) inhibitor (N-acetyl-L-cysteine; top) and an ROS inducer (Pyocyanin; bottom). (b) Representative image from patterned cardiomyocyte culture stained with an oxidative stress detection reagent (green) following treatment with an ROS inhibitor (top) and an ROS inducer (bottom). (c) Representative image from patterned cardiomyocyte culture stained with a nitric oxide (NO) detection reagent (blue) following treatment with an NO scavenger (c-PTIO; top) and an NO inducer (L-arginine; bottom). (d) Quantification of pixel intensity in images collected from cardiac cultures stained with a superoxide detection reagent. Samples analyzed were patterned cells treated with an ROS inhibitor (ROS-), patterned cells treated with an ROS inducer (ROS+), untreated unpatterned cells, and untreated patterned cells. Superoxide presence was significantly lower than all other groups in the ROS- samples, while it was significantly higher than all other groups examined in the ROS+ samples. No difference was observed between patterned and unpatterned samples. (e) Quantification of pixel intensity in images collected from cardiac cultures stained with an oxidative stress detection reagent. Samples analyzed were patterned cells treated with an ROS inhibitor (ROS-), patterned cells treated with an ROS inducer (ROS+), untreated unpatterned cells, and untreated patterned cells. ROS+ exhibited significantly higher values then all other groups, while ROS- was significantly lower than all other groups examined, with the exception of the patterned samples. No significant difference was observed between patterned and unpatterned samples. (f) Quantification of pixel intensity in images collected from cardiac cultures stained with an NO detection reagent. Samples analyzed were patterned cells treated with an NO scavenger (NO-), patterned cells treated with an NO inducer (NO+), untreated unpatterned cells, and untreated patterned cells. NO- was significantly lower than all other groups, while NO+ was significantly higher than all other groups examined. A significant difference was also observed between patterned and unpatterned samples. In all presented data, *p < 0.05.

**Supplementary Figure 5:**
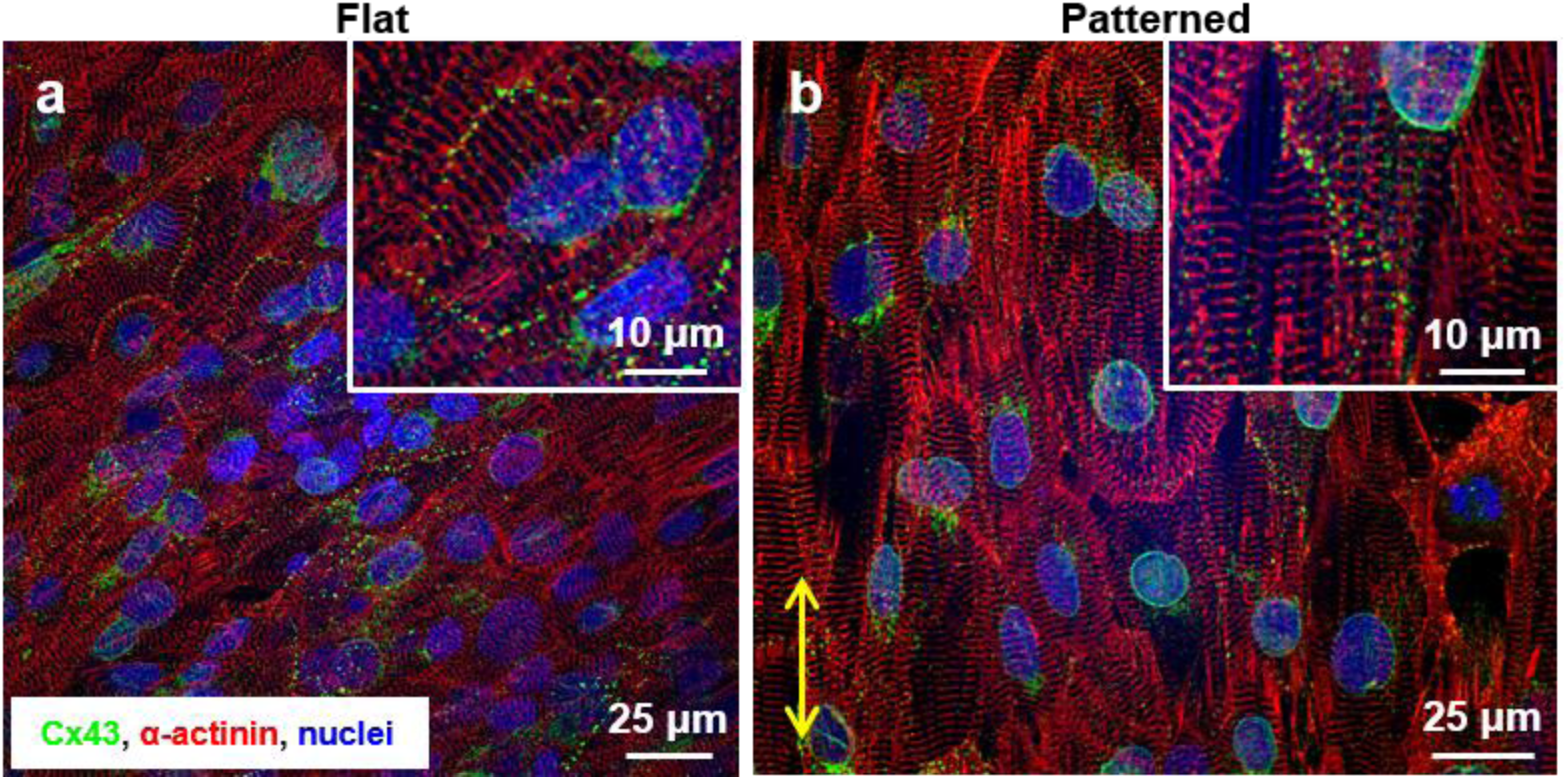
**Effect of nanotopography on spatial distribution of connexin 43 proteins in cultured hPSC-CMs.** (a) Representative immunostained image of hPSC-CMs on flat Nafion substrates showing random distribution of gap junctions. (b) Representative immunostained image of hPSC-CMs on patterned Nafion substrates showing more polar orientation of gap junctions. Yellow arrow indicates nanotopography orientation.

**Supplementary Figure 6:**
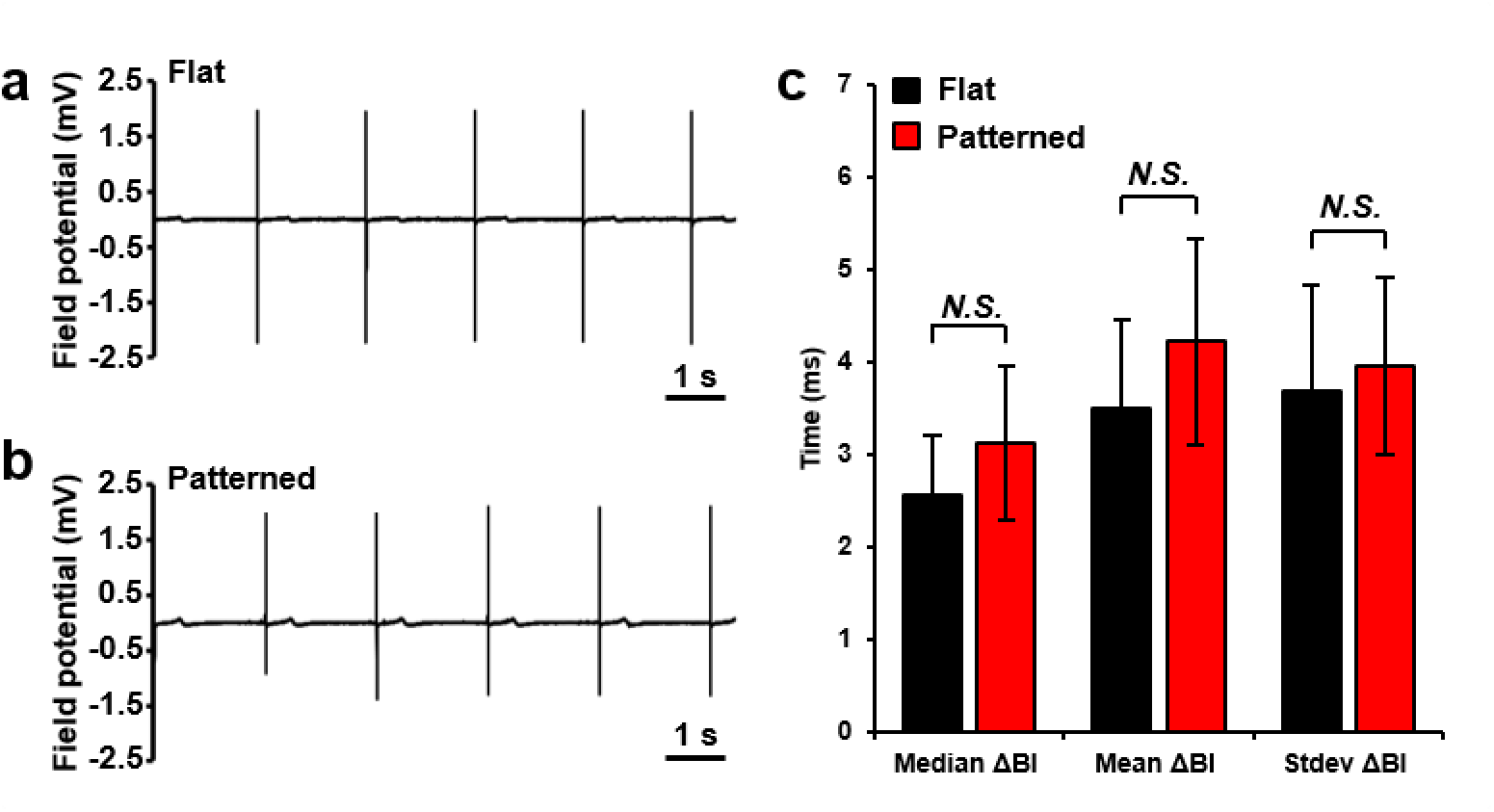
**Baseline electrophysiology in patterned and unpatterned hPSC-CM cultures.** (a) Representative baseline field potential recording from hPSC-CM monolayers maintained on flat Nafion MEAs for 21 days. (b) Representative baseline field potential recording from hPSC-CM monolayers maintained on patterned Nafion MEAs for 21 days. (c) Comparison of beat interval variability metrics calculated from analysis of hPSC-CMs maintained on patterned and unpatterned MEAs for 21 days. Median difference in beat interval (ΔBI), mean ΔBI, and the standard deviation of the ΔBI were compared in order to provide quantification of baseline arrhythmic properties in line with previously published methods^29^.

**Supplementary Figure 7:**
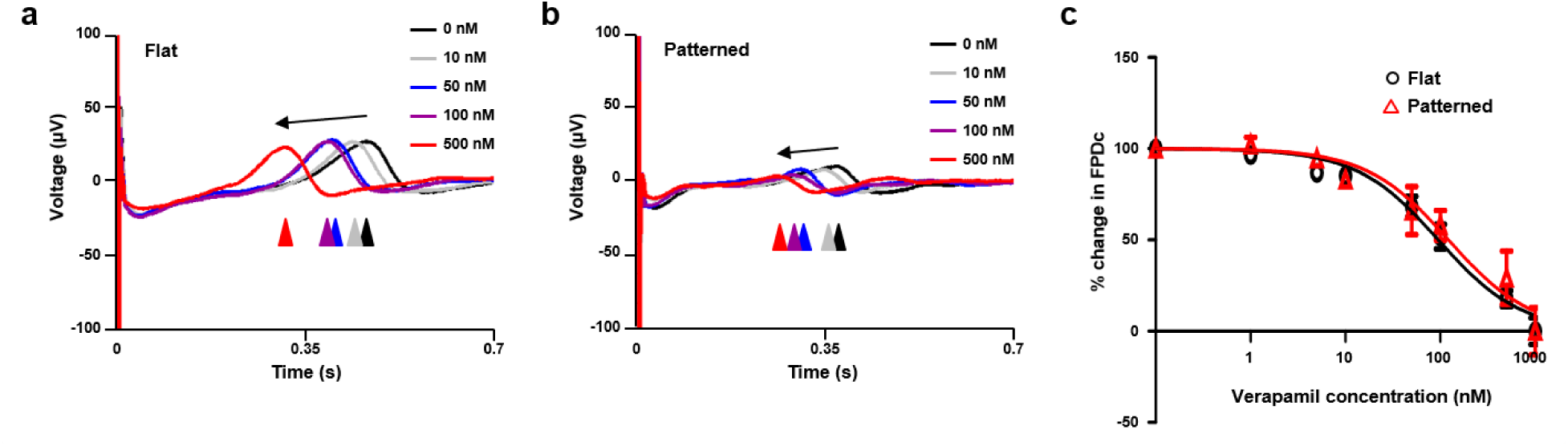
**Electrophysiological response of patterned and unpatterned hPSC-CMs to treatment with verapamil.** (a) Representative traces (averaged across 10 beats) recorded from hPSC-CM monolayers on flat MEAs and subjected to increasing doses of verapamil. (b) Representative traces recorded from hPSC-CM monolayers on nanoMEAs and subjected to increasing doses of verapamil. (c) Normalized dose response curve illustrating the effect for increasing concentrations of verapamil on the field potential durations corrected for beat rate (FPDc) of unpatterned and patterned hPSC-CMs. The R^2^ values for the unpatterned and patterned cultures were 0.86 and 0.60, respectively.

**Supplementary Table 1:** **Impedance measurements collected from bare and Nafion coated MEAs.** Impedance measurements were sampled at 1 kHz. n = 8.

| Condition | Before PEDOT (MΩ) | After PEDOT (MΩ) |
| --- | --- | --- |
| Bare | 2.56 ± 0.58 | 0.013 ± 0.00027 |
| Flat Nafion | 1.57 ± 0.16 | 0.010 ± 0.00054 |
| Patterned Nafion | 2.41 ± 0.20 | 0.015 ± 0.0019 |

## Supplementary Video Legends

**Supplementary Video 1: Real-time propagation map from hPSC-CMs maintained for 21 days on flat MEAs.** Spontaneous activation of the cardiac monolayer was observed consistently from the bottom right corner of the electrode area in this example.

**Supplementary Video 2: Real-time propagation map from hPSC-CMs maintained for 21 days on nanotopographically-patterned MEAs.** Nanotopographic patterns were aligned vertically in all patterned studies to ensure consistency of data. Spontaneous activation of the cardiac monolayer was observed consistently from the right of the electrode area in this example.

**Supplementary Video 3: Real-time propagation map from nanotopographically-patterned hPSC-CMs following 21 days *in vitro* and just prior to treatment with 500 nM carbenoxolone.** Spontaneous activation of the cardiac monolayer was observed consistently from the bottom right corner of the electrode area in this example.

**Supplementary Video 4: Real-time propagation map from nanotopographically-patterned hPSC-CMs following 21 days *in vitro* and 30 minutes after treatment with 500 nM carbenoxolone.** Spontaneous activation of the cardiac monolayer was observed consistently from the bottom right corner of the electrode area in this example.

